# Cross-reactive serum and memory B cell responses to spike protein in SARS-CoV-2 and endemic coronavirus infection

**DOI:** 10.1101/2020.09.22.308965

**Authors:** Ge Song, Wan-ting He, Sean Callaghan, Fabio Anzanello, Deli Huang, James Ricketts, Jonathan L. Torres, Nathan Beutler, Linghang Peng, Sirena Vargas, Jon Cassell, Mara Parren, Linlin Yang, Caroline Ignacio, Davey M. Smith, James E. Voss, David Nemazee, Andrew B Ward, Thomas Rogers, Dennis R. Burton, Raiees Andrabi

## Abstract

Pre-existing immune responses to seasonal endemic coronaviruses could have profound consequences for antibody responses to SARS-CoV-2, either induced in natural infection or through vaccination. Such consequences are well established in the influenza and flavivirus fields. A first step to establish whether pre-existing responses can impact SARS-CoV-2 infection is to understand the nature and extent of cross-reactivity in humans to coronaviruses. We compared serum antibody and memory B cell responses to coronavirus spike (S) proteins from pre-pandemic and SARS-CoV-2 convalescent donors using a series of binding and functional assays. We found weak evidence of pre-existing SARS-CoV-2 cross-reactive serum antibodies in pre-pandemic donors. However, we found stronger evidence of pre-existing cross-reactive memory B cells that were activated on SARS-CoV-2 infection. Monoclonal antibodies (mAbs) isolated from the donors showed varying degrees of cross-reactivity with betacoronaviruses, including SARS and endemic coronaviruses. None of the cross-reactive mAbs were neutralizing except for one that targeted the S2 subunit of the S protein. The results suggest that pre-existing immunity to endemic coronaviruses should be considered in evaluating antibody responses to SARS-CoV-2.

## Results and discussion

Well-known examples of pre-existing immunity to viruses influencing antibody (Ab) responses to related viruses include original antigenic sin (OAS) in influenza virus infections and antibody-dependent enhancement (ADE) in flavivirus infections ^1-3^. There is considerable interest in establishing whether Ab or T cell responses to SARS-CoV-2, through infection or vaccination, might be impacted by pre-existing immunity to other coronaviruses, particularly the endemic coronaviruses (endemic HCoVs), namely the betacoronaviruses (β-HCoV), HCoV-HKU1 and HCoV-OC43, and the alphacoronaviruses (α-HCoV), HCoV-NL63 and HCoV-229E, which are responsible for non-severe infections such as common colds ^4-8^. In principle, pre-existing immune perturbation effects could occur by interaction of SARS-CoV-2 with cross-reactive circulating serum Abs or with B cells bearing cross-reactive B cell receptors (BCRs) or T cells with cross-reactive T cell receptors (TCRs). While a number of studies have reported on cross-reactive T cells and serum Abs ^6,8-12^, we investigate here both Ab and BCR cross-reactivities.

Since individuals who have been infected with SARS-CoV-2 will generally also have been infected with endemic HCoVs, we chose to compare COVID-19 and pre-pandemic donors in terms of serum Abs and BCRs with specificity for the spike (S) protein. The rationale was that the pre-pandemic donor cross-reactive responses could only be due to endemic HCoV infection. However, the COVID-19 cohort could reveal the effects of SARS-CoV-2 infection on cross-reactive responses.

To assess serum Ab S-protein binding in the two cohorts, we used cell-surface and recombinant soluble S proteins. First, we developed and utilized a high-throughput flow cytometry-based cell surface spike binding assay (Cell-based ELISA; CELISA). COVID-19 convalescent sera from 36 donors showed strong reactivity to the SARS-CoV-2 spike in the vast majority of infected donors (Fig. 1a, supplementary Fig. 1), somewhat lower reactivity with the SARS-CoV-1 spike and much lower reactivity with the MERS-CoV spike in a pattern consistent with sequence conservation between the 3 viruses. COVID sera also exhibited strong cross-reactivity with endemic HCoV spikes, especially with the HCoV-HKU1 and HCoV-OC43 β-HCoVs (Fig. 1a). The α-HCoV-derived HCoV-NL63 spike was least reactive among the 4 endemic HCoVs. Next, we tested sera from a cohort of 36 healthy human donors whose samples were collected pre-pandemic. The sera showed minimal or no reactivity to SARS-CoV-2/CoV-1 and MERS-CoV spikes but showed strong binding to the endemic HCoV spikes, especially against the HCoV-HKU1 and HCoV-OC43 β-HCoVs (Fig. 1, supplementary Fig. 1). The results suggest that the pre-pandemic sera, at least in our cohort, possess low levels of pre-existing SARS-CoV-2 circulating Abs.

**Fig. 1.**
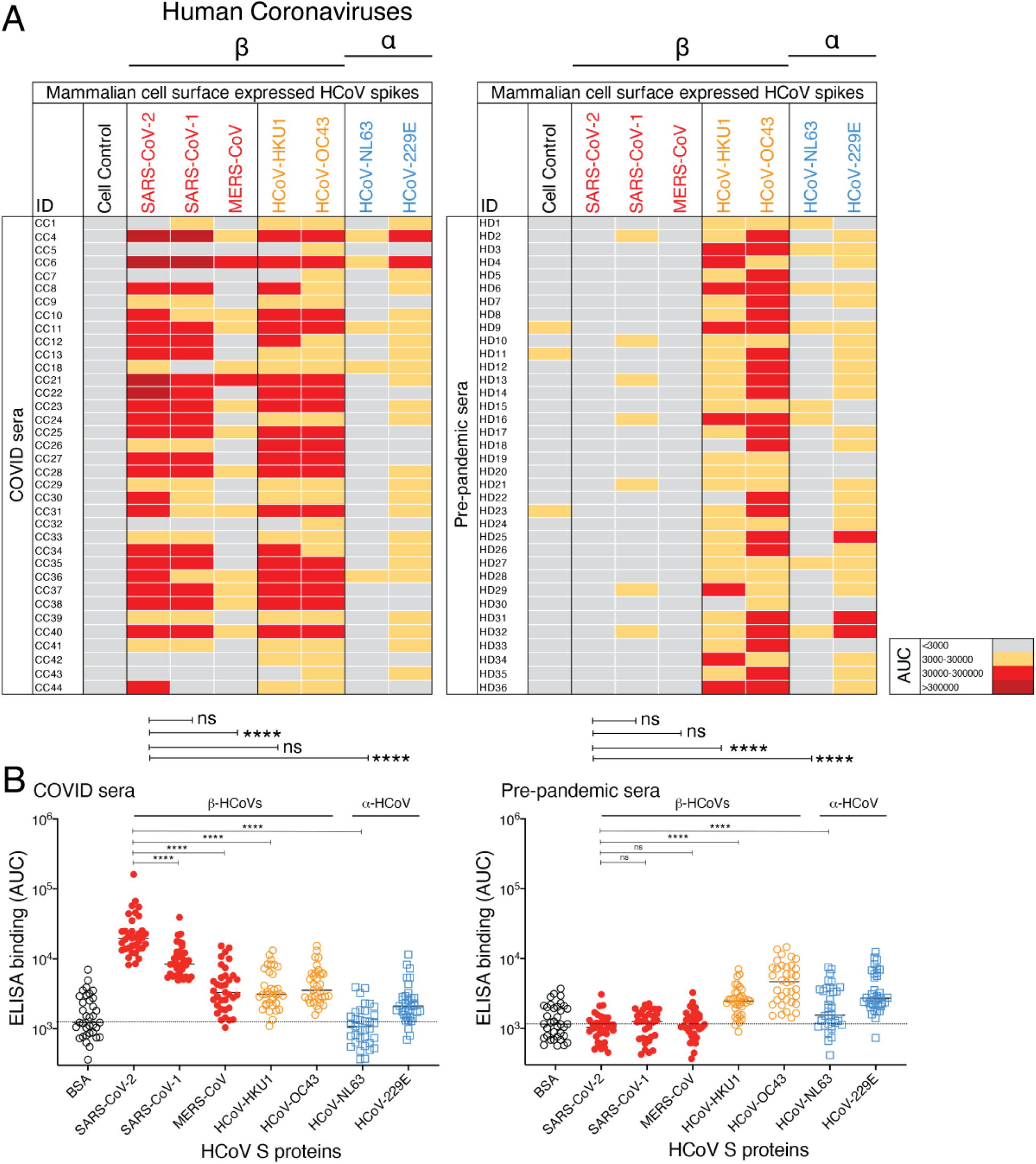
Reactivity of COVID and pre-pandemic human sera with cell surface-expressed human coronaviruses spikes and their soluble S-protein versions. **A**. Heatmap showing cell-based flow cytometry binding (CELISA) of COVID and pre-pandemic donor sera with 293T cell surface-expressed full-length spike proteins from β-(SARS-CoV-2, SARS-CoV-1, MERS-CoV, HCoV-HKU1, HCoV-OC43) and α-(HCoV-NL63 and HCoV-229E) human coronaviruses (HCoVs). Sera were titrated (6 dilutions-starting at 1:30 dilution) and the extent of binding to cell surface-expressed HCoVs was recorded by % positive cells, as detected by PE-conjugated anti-human-Fc secondary Ab using flow cytometry. Area-under-the-curve (AUC) was calculated for each binding titration curve and the antibody titer levels are color-coded as indicated in the key. Binding of sera to vector-only plasmid (non-spike) transfected 293T cells served as a control for non-specific binding. **B**. ELISA binding of COVID and pre-pandemic donor sera to soluble S-proteins from β-(SARS-CoV-2, SARS-CoV-1, MERS-CoV, HCoV-HKU1, HCoV-OC43) and α-(HCoV-NL63 and HCoV-229E) HCoVs. Serum dilutions (8 dilutions-starting at 1:30 dilution) were titrated against the S-proteins and the binding was detected as OD405 absorbance. AUC representing the extent of binding was calculated from binding curves of COVID (left) and pre-pandemic (right) sera with S-proteins and comparisons of antibody binding titers are shown. Binding to BSA served as a control for non-specific binding by the sera. Statistical comparisons between two groups were performed using a Mann-Whitney test, (**p <0.01; ***p < 0.001, ****p < 0.0001; ns-p >0.05).

To further investigate, we generated recombinant soluble S proteins of all 7 HCoVs using a general stabilization strategy described elsewhere ^13-15^. ELISA showed a similar binding pattern of the COVID and pre-pandemic sera as the CELISA (Fig. 1B, supplementary Fig. 1). The SARS-CoV-2 S specific binding of COVID sera in the two assay formats (CELISA versus ELISA) correlated strongly (r = 0.92, p < 0.001) (supplementary Fig. 2), CELISA being more sensitive overall. We also tested the neutralization of the COVID sera with SARS-CoV-2 and the ID_50_ neutralization titers positively correlated with both binding assays (CELISA (r = 0.72, p < 0.0001), ELISA (r = 0.68, p < 0.0001)) (supplementary Fig. 2). Overall, both CELISA and ELISA revealed binding Abs to all 7 HCoV spikes in COVID sera but only to endemic HCoVs in the pre-pandemic sera.

**Fig. 2.**
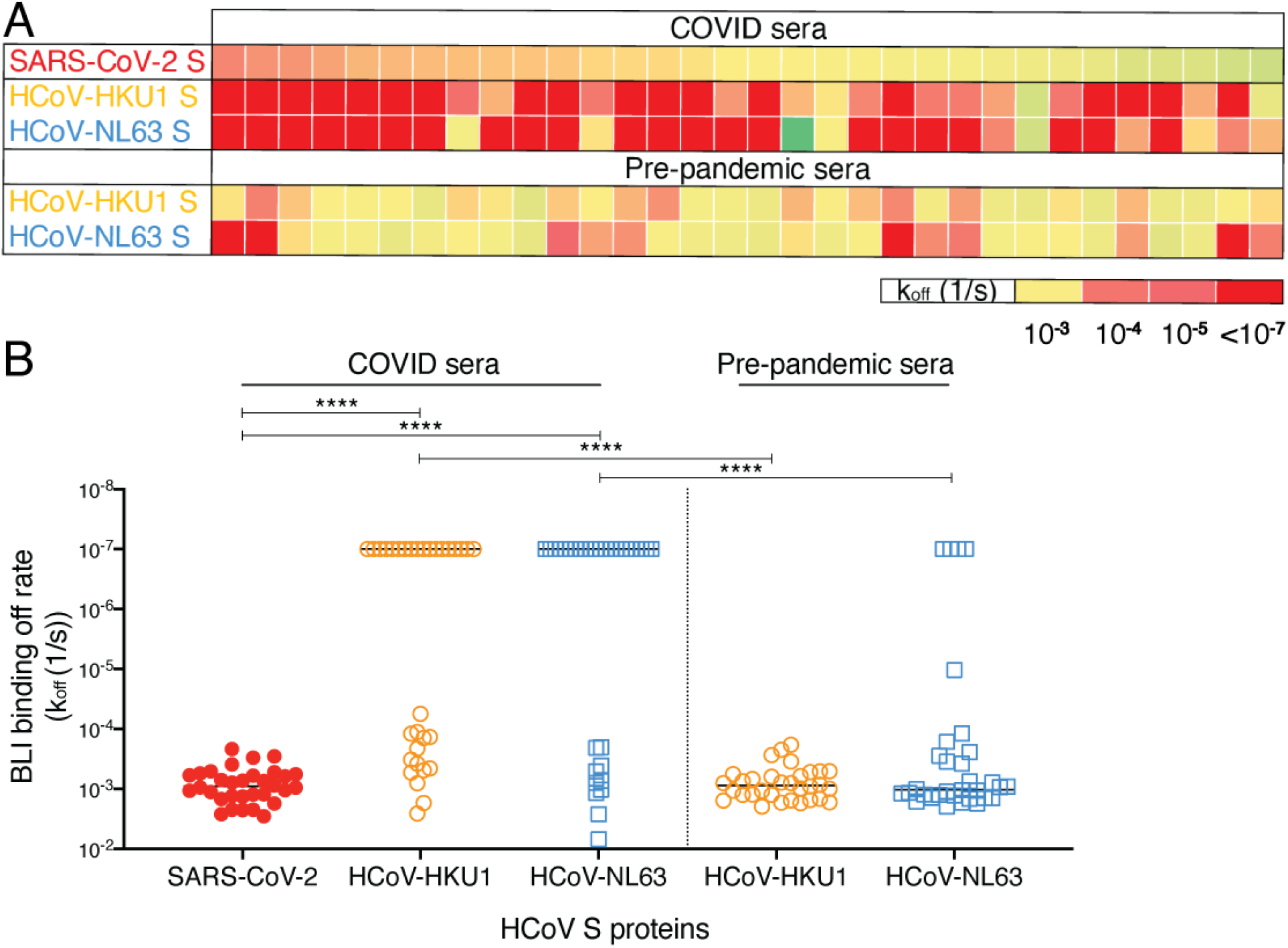
BioLayer Interferometry binding of COVID and pre-pandemic serum antibodies to SARS-CoV-2 and endemic HCoV S-proteins. **A**. Heatmap summarizing the apparent BLI binding off-rates (k_off_ (1/s)) of the COVID and pre-pandemic human serum antibodies to SARS-CoV-2 S and endemic β-HCoV, HCoV-HKU1 and α-HCoV, HCoV-NL63 S-proteins. Biotinylated HCoV S-proteins (100nM) were captured on streptavidin biosensors to achieve binding of at least 1 response unit. The S-protein-immobilized biosensors were immersed in 1:40 serum dilution solution with serum antibodies as the analyte and the association (120 s; 180-300) and dissociation (240 s; 300-540) steps were conducted to detect the kinetics of antibody-protein interaction. k_off_ (1/s) dissociation rates for each antibody-antigen interaction are shown. **B**. Off-rates for binding of serum antibodies from COVID donors and from pre-pandemic donors to SARS-CoV-2 S and endemic HCoV, HCoV-HKU1 and HCoV-NL63, S proteins. Significantly lower dissociation off-rates are observed for COVID compared to pre-pandemic sera. Statistical comparisons between the two groups were performed using a Mann-Whitney test.

To assess whether SARS-CoV-2 infection may impact serum Ab titers to endemic HCoVs, we compared Ab titers to endemic HCoV S-protein in sera from COVID and pre-pandemic cohorts. Higher CELISA Ab titers to endemic HCoV-HKU1 S-protein, but not for other HCoV spikes (HCoV-OC43, HCoV-NL63 and HCoV-229E) were observed in the COVID cohort compared to the pre-pandemic cohort (supplementary Fig. 3). The result suggests that SARS-CoV-2 infection may boost titers to the related HCoV-HKU1 spike ^16,17^. To further investigate, we divided individuals from the COVID cohort into two groups, one with the higher SARS-CoV-2 spike Ab titers (AUC > 85,000) and the other with lower titers (AUC < 85,000). Consistent with the above result, the COVID sera with higher SARS-CoV-2 titers showed significantly higher binding to HCoV-HKU1 and HCoV-OC43 S-proteins compared to the low titer group (supplementary Fig. 3). The α-HCoVs HCoV-NL63 and HCoV-229E spike binding antibody titers were comparable between the two groups and served as a control (supplementary Fig. 3). Since the two cohorts are not matched in terms of a number of parameters and are of limited size, any conclusions should be treated with caution. Nevertheless, it is noteworthy that SARS-CoV-2 infection is apparently associated with enhanced β-HCoVs S-protein Ab responses. A key question is whether the enhanced responses arise from de novo B cell responses or from a recall response of B cells originally activated by an endemic HCoV virus infection.

**Fig. 3.**
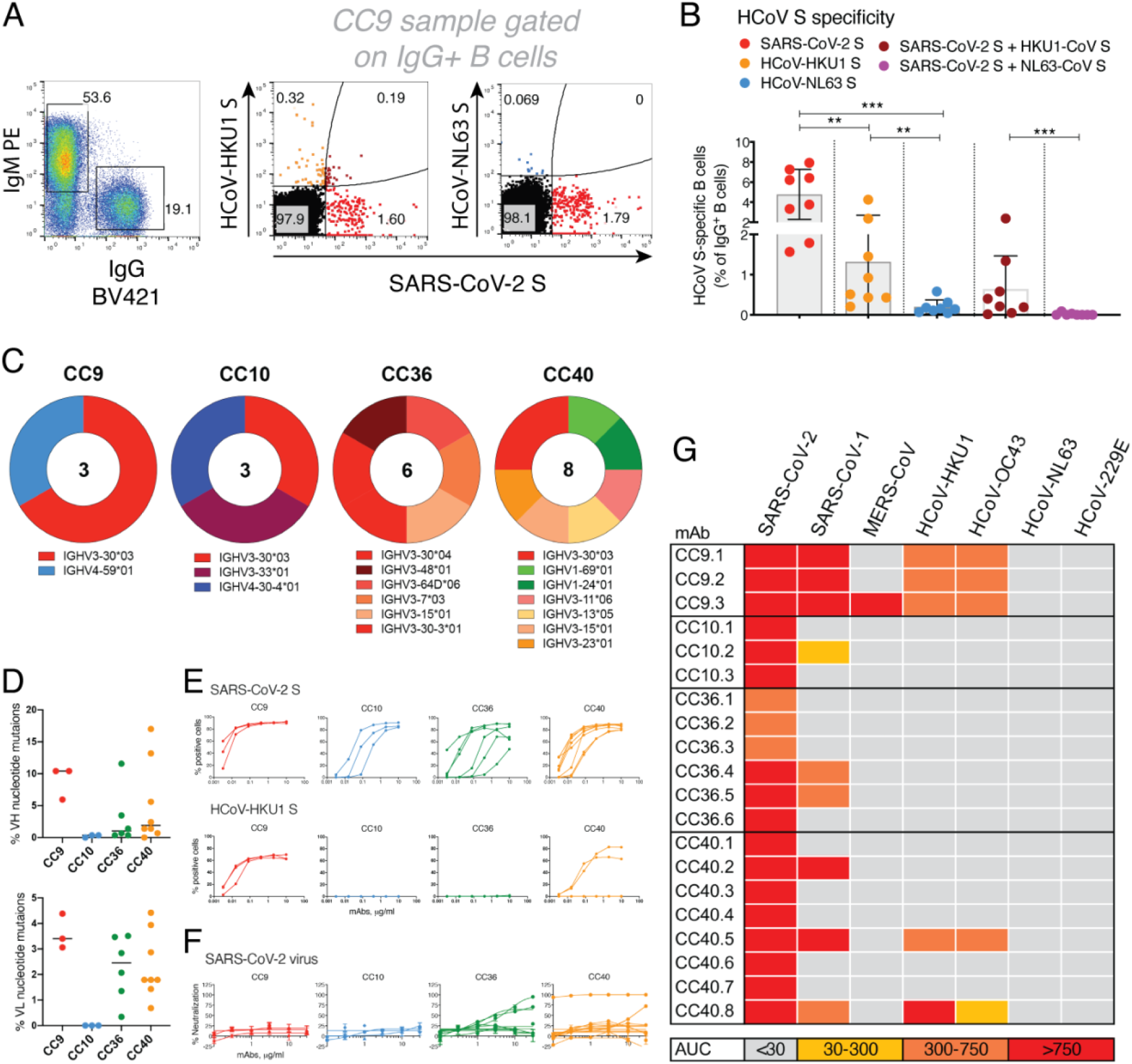
SARS-CoV-2 S and endemic HCoV S-protein specific cross-reactive IgG+ memory B cells from COVID donors and isolation of and characterization of mAbs. **A-B**. Flow cytometry analysis showing the single B cell sorting strategy for COVID representative donor CC9 and frequencies of SARS-CoV-2 S and endemic β-HCoV, HCoV-HKU1 and α-HCoV, HCoV-NL63 S-protein specific memory B cells in 8 select COVID donors. The B cells were gated as SSL, CD4-, CD8-, CD11C-, IgD-, IgM-, CD19+, IgG+. The frequencies of HCoV S-protein-specific IgG memory B cells were as follows; SARS-CoV-2 S (up to ∼8% - range = ∼1.6-8%), HCoV-HKU1 S (up to ∼4.3% - range = ∼0.2-4.3%), HCoV-NL63 S (up to ∼0.6% - range = ∼0.04-0.6%) protein single positive and SARS-CoV-2/HCoV-HKU1 S (up to ∼2.4% - range = ∼0.02-2.4%) and SARS-CoV-2/HCoV-NL63 S-protein (up to ∼0.09% - range = ∼0-0.09%) double positives. SARS-CoV-2 infected donors showed the presence of SARS-CoV-2/HCoV-HKU1 S-protein cross-reactive IgG memory B cells. A Mann-Whitney test was used to compare the levels of HCoV S-protein specific IgG memory B cells and the p-values for each comparison are indicated. **p <0.01; ***p < 0.001. **C**. Pie plots showing immunoglobulin heavy chain distribution of mAbs isolated from 4 COVID donors, CC9, CC10, CC36 and CC40. The majority of the mAbs were encoded by the IgVH3 immunoglobulin gene family. **D**. Plots showing % nucleotide mutations in heavy (VH) and light (VL) chains of isolated mAbs across different individuals. The VH and VL mutations ranged from 0-17% and 0-4.5%, respectively. **E**. CELISA binding curves of isolated mAbs from 4 COVID donors with SARS-CoV-2 and HCoV-HKU1 spikes expressed on 293T cells. Binding to HCoV spikes is recorded as % positive cells using a flow cytometry method. 5 mAbs, 3 from the CC9 donor and 2 from the CC40 donor show cross-reactive binding to SARS-CoV-2 and HCoV-HKU1 spikes. **F**. Neutralization of SARS-CoV-2 by mAbs isolated from COVID donors. 4 mAbs, 2 each from donors, CC36 and CC40, show neutralization of SARS-CoV-2. **G**. Heatmap showing CELISA binding of COVID mAbs to 7 HCoV spikes. Binding represented as area-under-the-curve (AUC) is derived from CELISA binding titrations of mAbs with cell surface-expressed HCoV spikes and the extent of binding is color-coded. 5 mAbs show cross-reactive binding across β-HCoV spikes.

We were encouraged to look more closely at the Abs involved by Bio-Layer Interferometry (BLI). Polyclonal serum antibodies were used as analytes with biotinylated S proteins captured on streptavidin biosensors. Since the concentrations of the S protein specific polyclonal Abs in the sera are unknown, these measurements can provide an estimate of antibody dissociation off-rates (k_off_, which is antibody concentration independent) but not binding constants ^18^. Slower dissociation off-rates would indicate greater affinity maturation of antibodies with a given S protein ^19^. It is important to note that the off-rates are likely associated with bivalent IgG binding (avidity) in the format used. Consistent with the notion of SARS-CoV-2 infection activating a recall of cross-reactive HCoV S specific Abs, the COVID sera Abs exhibited significantly slower off-rates with HCoV-HKU1 and HCoV-NL63 S-proteins compared to pre-pandemic sera Abs (Fig. 2A-B, supplementary Fig. 4).

**Fig. 4.**
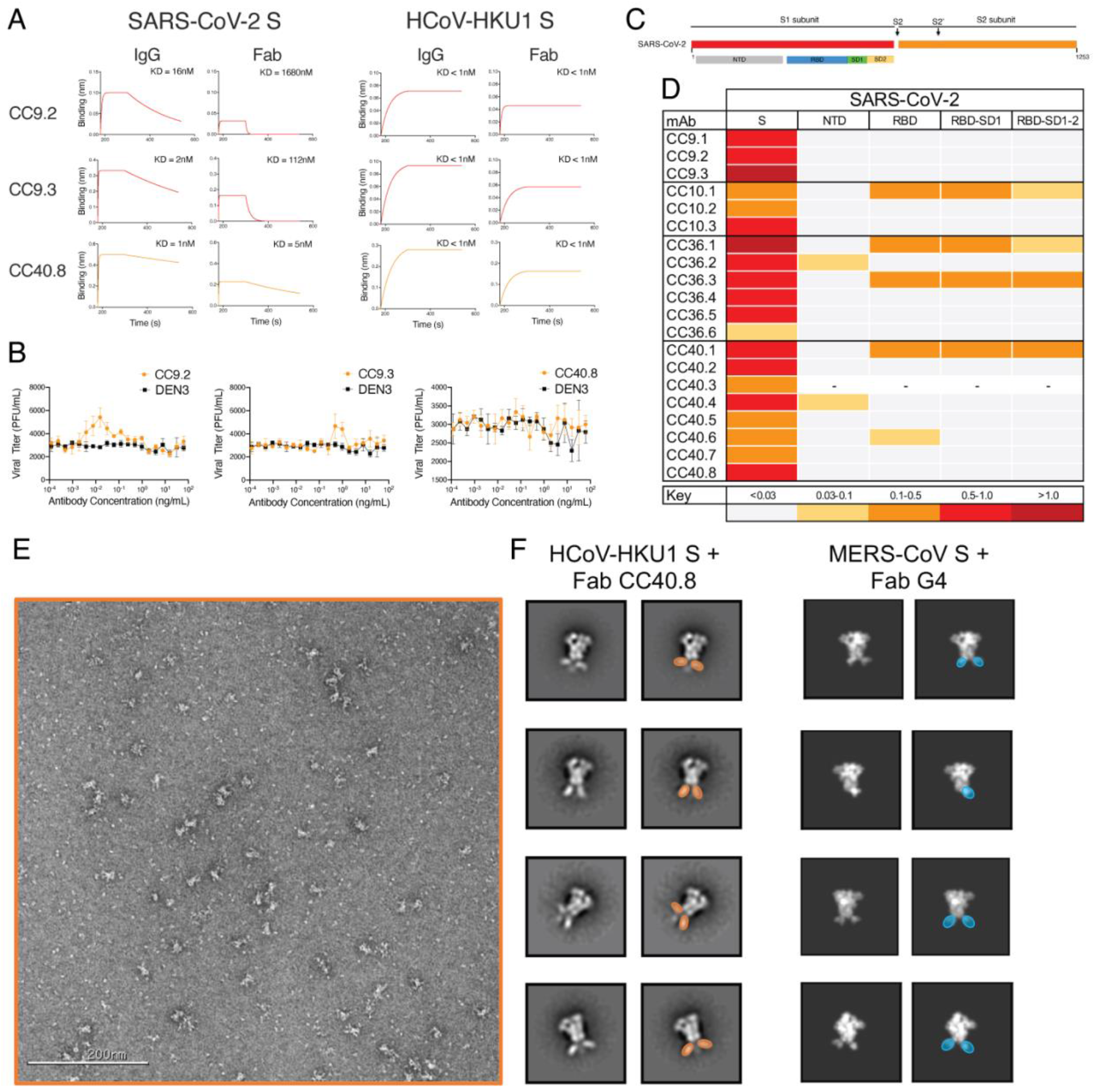
Binding, ADE and epitope specificities of SARS-CoV-2/HCoV-HKU1 S-protein specific cross-reactive mAbs. **A**. BLI of SARS-CoV-2 and HCoV-HKU1 S-protein-specific cross-reactive mAbs. BLI binding of both IgG and Fab versions of 3 cross-reactive mAbs (CC9.2, CC9.3 and CC40.8) to SARS-CoV-2 and HCoV-HKU1 S-proteins was tested and the binding curves show association (120 s; 180-300) and dissociation rates (240 s; 300-540). BLI binding of antibody-S-protein combinations shows more stable binding (higher binding constants (KDs)) of cross-reactive mAbs HCoV-HKU1 compared to the SARS-CoV-2 S protein. **B**. Antibody Dependent Enhancement (ADE) activities of cross-reactive mAbs, CC9.2, CC9.3 and CC40.8 bonding to SARS-CoV-2 live virus using FcγRIIa (K562) and FcγRIIb (Daudi)-expressing target cells. A dengue antibody, DEN3, was used as a control. **C-D**. Epitope mapping of the mAbs binding to domains and subdomains of SARS-CoV-2 S-protein, NTD, RBD, RBD-SD1 and RBD-SD1-2 and heatmap showing BLI responses for each protein. The extent of binding responses is color coded. 5 mAbs were specific for RBD, 2 for NTD and the remaining mAbs displayed binding only to the whole S protein. **E-F**. Negative stain electron microscopy of HCoV-HKU1 S-protein + Fab CC40.8 complex and comparison to MERS-CoV S + Fab G4 complex. (E) Raw micrograph of HCoV-HKU1 S in complex with Fab CC40.8. (F) Select reference-free 2D class averages with Fabs colored in orange for Fab CC40.8 and blue for Fab G4, which in 2D appear to bind a proximal epitope at the base of the trimer. 2D projections for MERS-CoV S-protein in complex with Fab G4 were generated in EMAN2 from PDB 5W9J.

Having probed serum cross-reactivity between coronaviruses, we next investigated memory B cells in COVID individuals. We examined the reactivities of IgG+ memory B cells in 8 select COVID donors (based on differential binding to HCoV spikes (Fig. 1) with SARS-CoV-2, HCoV-HKU1 (β-HCoV) and HCoV-NL63 (α-HCoV) S-proteins by flow cytometry. Up to ∼8% SARS-CoV-2 S-protein, ∼4.3% HCoV-HKU1 S-protein and ∼0.6% for HCoV-NL63 S-protein-specific B cells were identified (Fig. 3B) in a frequency pattern consistent with serum antibody binding titers.

To probe the specificities of SARS-CoV-2/endemic HCoV cross-reactive Abs, we sorted single B cells for either SARS-CoV-2/HCoV-HKU-1 or SARS-CoV-2/HCoV-NL63 CoV S-protein double positivity. We isolated 20 S-protein-specific mAbs from 4 COVID donors, CC9 (n=3), CC10 (n=3), CC36 (n=6) and CC40 (n=8) (Fig. 3C, supplementary Fig. 5) but only 5 mAbs, 3 from the CC9 donor and 2 from the CC40 donor, exhibited cross-reactive binding with HCoV-HKU1 spike (Fig. 3E). Two of the cross-reactive mAbs from the CC9 donor (CC9.1 and CC9.2) were clonally related. All 5 of the SARS-CoV-2/ HCoV-HKU-1 cross-reactive mAbs displayed binding to the genetically related β-HCoV, HCoV-OC43, spike but not to the α-HCoVs, HCoV-NL63 and HCoV-229E, spikes (Fig. 3G, supplementary Fig. 6). Notably, one mAb (CC9.3) exhibited binding to 5 out of the 7 HCoVs, including the MERS-CoV S-protein (Fig. 3G, supplementary Fig. 6) suggesting targeting of a highly conserved epitope on β-HCoV spikes. One of the 4 SARS-CoV-2/HKU1-CoV S cross-reactive mAbs (CC40.8) showed weak cross neutralization against SARS-CoV-2 and SARS-CoV-1 viruses (supplementary Fig. 6). Except for CC9.3 mAb, all cross-reactive mAbs were encoded by VH3 family gene heavy chains (supplementary Figs. 5 and 6) and possessed 5.6-13.2% (median = 10.4%) VH and 3.1-4.4% (median = 3.9%) VL nucleotide SHMs (Fig. 3D supplementary Fig. 5).

In principle, the SARS-CoV-2/HCOV-HKU1 S cross-reactive memory B cells could be pre-existing in the COVID donors and show cross-reactivity with SARS-CoV-2 or originate from the SARS-CoV-2 infection and show cross-reactivity with HCoV-HKU1 S protein. The levels of SHM in the 5 cross-reactive mAbs listed above argue for the first explanation. To gain further insight, we conducted BLI binding studies on the 3 cross-reactive mAbs, CC9.2, CC9.3 and CC40.8 (Fig. 4A). Both bivalent IgGs and monovalent Fabs showed enhanced binding affinity to HCoV-HKU1 S-protein compared to SARS-CoV-2 S-protein (Fig. 4A) again consistent with the notion that the Abs (BCRs) arise from a pre-existing HCoV-HKU1 S response. The serum and BCR data are then consistent. The data above suggests elevated serum levels of Abs to HCoV-HKU1 S-protein in COVID donors compared to pre-pandemic donors (Fig. 2A-B) is consistent with the notion that SARS-CoV-2 activates B cells expressing pre-existing HCoV-HKU1 S-protein specific BCRs to secrete the corresponding Abs.

One mechanism by which pre-existing cross-reactive antibodies might influence the course of SARS-CoV-2 infection is ADE. Therefore, we investigated potential ADE of the 3 cross-reactive Abs using a SARS-CoV-2 live virus assay (Fig. 4B). Of the 3 cross-reactive antibodies, CC9.3 mAb showed a marginal increase (2-fold) in infection of SARS-CoV-2 virus in the FcγRIIa (K562) and FcγRIIb (Daudi) expressing target cells that can mediate ADE. Further *in vivo* assessment would be needed to determine if this activity is associated with any meaningful physiological effects.

To map the epitope specificities of the cross-reactive mAbs, we evaluated binding to a number of fragments of the S-protein (Fig. 4C-D). Notably, all 5 of the SARS-CoV-2/HKU1-CoV cross-reactive mAbs failed to bind any of the S1 subunit domains or subdomains, suggesting targeting to the more conserved S2 subunit. To identify the cross-reactive neutralizing epitope recognized by mAb CC40.8, we conducted structural studies of the antibody with the HKU1-CoV S protein. Using single particle negative stain electron microscopy (nsEM) we observed that CC40.8 bound to the HCoV-HKU1 S trimer near the bottom of the S2 domain (Fig. 4E-F). The Fab density in the 2D class averages was blurry suggesting binding to a flexible surface exposed peptide. The flexibility also precluded further 3D reconstruction.

Despite the requirement of double positivity in the B cell sorting, 15/20 mAbs were largely specific for SARS-CoV-2. Again, like cross-reactive mAbs above, the vast majority of SARS-CoV-2 specific mAbs were encoded by VH3 family gene-encoded heavy chains (Fig. 3C, supplementary Fig. 5), consistent with other studies ^20-26^. Compared to the cross-reactive mAbs, the nucleotide SHM levels in SARS-CoV-2 specific mAbs were much lower (VH, 0-17% (median = 0.7%) VL, 0-3.5% (median = 1.8%)) (Fig. 3D supplementary Fig. 5). 3 of the 15 SARS-CoV-2 S specific mAbs showed neutralization against SARS-CoV-2 virus, CC40.1 being the most potent (Fig. 3F, supplementary Fig. 6). Some of the SARS-CoV-2 specific mAbs exhibited cross-reactive binding with SARS-CoV-1 S protein but none neutralized SARS-CoV-1.

In conclusion, using a range of immune monitoring assays, we compared the serum and memory B cell responses to the S-protein from all 7 coronaviruses infecting humans in SARS-CoV-2 donors and in pre-pandemic donors. In sera from our pre-pandemic cohort, we found no evidence of pre-existing SARS-CoV-2 S-protein reactive antibodies that resulted from endemic HCoV infections. A recent study has however reported the presence of SARS-CoV-2 S-protein reactive antibodies in a small fraction of pre-pandemic human sera ^11^. An in-depth examination for the presence of SARS-CoV-2 S-protein reactive antibodies in large pre-pandemic human cohorts is warranted to reliably determine the frequency of such antibodies. Notably, we observed serum levels of endemic HCoV S-protein antibodies were higher in SARS-CoV-2-experienced donors and memory B cell studies suggested these likely arose from SARS-CoV-2 infection activating cross-reactive endemic HCoV S-protein-specific B cells. Cross-reactive mAbs largely target the more conserved S2 subunit on S-proteins and we identified a SARS-CoV-2 cross-neutralizing epitope that could facilitate vaccine design and antibody-based intervention strategies. Indeed, studies have shown targeting of conserved S2 subunit neutralizing epitopes in SARS-CoV-2 infected donors and by SARS-CoV-1 nAbs that may potentially display activities against a broader range of human coronaviruses ^27-30^. Overall, our study highlights the need to understand fully the nature of pre-existing endemic HCoV immunity in large and diverse human cohorts as vaccination of hundreds of millions of people against COVID-19 is considered.

## Acknowledgements

We thank all the COVID-19 cohort and healthy human cohort participants for donating samples. This work was supported by the NIH CHAVD (UM1 AI44462 to A.B.W. and D.R.B.), and R01 (AI132317, AI073148 to D.N.) awards, the IAVI Neutralizing Antibody Center, the Bill and Melinda Gates Foundation (OPP 1170236 to A.B.W. and D.R.B.). This work was also supported by the John and Mary Tu Foundation and the Pendleton Trust.

## Author contributions

R.A. and D.R.B. conceived and designed the study. T.F.R., N.B., J.R., M.P., L.Y., C.I. and D.M.S. recruited donors, collected and processed plasma and PBMC samples; G.S., W.H., S.C., F.A., D.H., J.R., J.L.T., N.B., L.P., S.V., and J.C. made substantial contributions to the acquisition of data and data analyses; G.S., W.H., S.C., F.A., D.H., J.R., J.L.T., N.B., L.P., S.V., J.C., J.E.V., D.N., A.B.W., T.F.R., D.R.B., and R.A. designed experiments and analyzed the data. R.A. and D.R.B. wrote the paper and all authors reviewed and edited the paper.

## Methods

### Plasmid construction for full-length and recombinant soluble proteins

To generate full-length human coronavirus plasmids, the spike genes were synthesized by GeneArt (Life Technologies). The SARS-CoV-1 (1255 amino acids; GenBank: AAP13567), SARS-CoV-2 (1273 amino acids; GenBank: MN908947), MERS-CoV (1353 amino acids; GenBank: APB87319.1), HCoV-HKU1 (1356 amino acids; GenBank: YP_173238.1), HCoV-OC43 (1361 amino acids; GenBank: AAX84792.1), HCoV-NL63 (1356 amino acids; GenBank: YP_003767.1) and HCoV-229E (1173 amino acids; GenBank: NP_073551.1) were cloned into the mammalian expression vector phCMV3 (Genlantis, USA) using PstI and BamH restriction sites. To express the soluble S ectodomain protein SARS-CoV-1 (residue 1-1190), SARS-CoV-2 (residue 1-1208), MERS-CoV (residue 1-1291), HCoV-HKU1 (residue 1-1295), HCoV-OC43 (residue 1-1300) and HCoV-NL63 (residue 1-1291), HCoV-229E (residue 1-1110), the corresponding DNA fragments were PCR amplified and constructed into vector phCMV3 using a Gibson assembly kit. To trimerize the soluble S proteins and stabilize them in the prefusion state, we incorporated a C-terminal T4 fibritin trimerization motif in the C-terminal of each constructs and two consecutive proline substitutions in the S2 subunit ^13-15^. To be specific, the K968/V969 in SARS-CoV-1, the K986/V987 in SARS-CoV-2, the V1060/L1061 in MERS-CoV, the A1071/L1072 in HCoV-HKU1, the A1078/L1079 in HCoV-OC43, the S1052/I1053 in HCoV-NL63 and the T871/I872 in HCoV-229E were replaced by proline residues. Additionally, the S2 cleavage sites in each protein were replaced with a “GSAS” linker peptide. To facilitate the purification and biotin labeling of the soluble protein, the HRV-3C protease cleavage site, 6X HisTag, and AviTag spaced by GS-linkers were added to the C-terminus of the constructs, as needed. To express the SARS-CoV-2 N-terminal domain-NTD (residue 1-290), receptor-binding domain-RBD (residue 320-527), RBD-SD1 (residue 320-591), and RBD-SD1-2 (residue 320-681) subdomains, we amplified the DNA fragments by PCR reaction using the SARS-CoV-2 plasmid as template. All the DNA fragments were cloned into the vector phCMV3 (Genlantis, USA) in frame with the original secretion signal or the Tissue Plasminogen Activator (TPA) leader sequence. All the truncation proteins were fused to the C-terminal 6X HisTag, and AviTag spaced by GS-linkers to aid protein purification and biotinylation.

### Expression and purification of the proteins

To express the soluble S ectodomain proteins of each human coronavirus and the truncated versions, the plasmids were transfected into FreeStyle293F cells (Thermo Fisher). For general production, 350 ug plasmids were transfected into 1L FreeStyle293F cells at the density of 1 million cells/mL. We mixed 350 ug plasmids with 16mL transfectagro™ (Corning) and 1.8 mL 40K PEI (1mg/mL) with 16mL transfectagro™ in separate 50 mL conical tubes. We filtered the plasmid mixture with 0.22 μm Steriflip™ Sterile Disposable Vacuum Filter Units (MilliporeSigma™) before combining it with the PEI mixture. After gently mixing the two components, the combined solution rested at room temperature for 30 min and was poured into 1 L FreeStyle293F cell culture. To harvest the soluble proteins, the cell cultures were centrifuged at 3500 rpm for 15 min on day 4 after transfection. The supernatants were filtered through the 0.22 μm membrane and stored in a glass bottle at 4 °C before purification. The His-tagged proteins were purified with the HisPur Ni-NTA Resin (Thermo Fisher). To eliminate nonspecific binding proteins, each column was washed with at least 3 bed volumes of wash buffer (25 mM Imidazole, pH 7.4). To elute the purified proteins from the column, we loaded 25 mL of the elution buffer (250 mM Imidazole, pH 7.4) at slow gravity speed (∼4 sec/drop). Proteins without His tags were purified with GNL columns (Vector Labs). The bound proteins were washed with PBS and then eluted with 50 mL of 1M Methyl α-D-mannopyranoside (Sigma M6882-500G) in PBS. By using Amicon tubes, we buffer exchanged the solution with PBS and concentrated the proteins. The proteins were further purified by size-exclusion chromatography using a Superdex 200 Increase 10/300 GL column (GE Healthcare). The selected fractions were pooled and concentrated again for further use.

### Biotinylation of proteins

Random biotinylation of S proteins was conducted using EZ-Link NHS-PEG Solid-Phase Biotinylation Kit (Thermo Scientific #21440). 10ul DMSO were added per tube for making concentrated biotin stock, 1ul of which were diluted into 170ul water before use. Coronavirus spike proteins were concentrated to 7-9 mg/ml using 100K Amicon tubes in PBS, then aliquoted into 30ul in PCR tubes. 3ul of the diluted biotin were added into each aliquot of concentrated protein and incubated on ice for 3h. After reaction, buffer exchange for the protein was performed using PBS to remove excess biotin. BirA biotinylation of S proteins was conducted using BirA biotin-protein ligase bulk reaction kit (Avidity). Coronavirus S proteins with Avi-tags were concentrated to 7-9 mg/ml using 100K Amicon tubes in TBS, then aliquoted into 50ul in PCR tubes. 7.5ul of BioB Mix, 7.5ul of Biotin200, and 5ul of BirA ligase (3mg/ml) were added per tube. The mixture was incubated on ice for 3h, followed by size-exclusion chromatography to segregate the biotinylated protein and the excess biotin. The extend of biotinylation was evaluated by BioLayer Interferometry binding value using streptavidin biosensors.

### CELISA binding

Binding of serum antibodies or mAbs to human coronavirus spike proteins expressed on HEK293T cell surface was determined by flow cytometry, as described previously ^31^. HEK293T cells were transfected with plasmids encoding full-length coronavirus spikes including SARS-CoV-1, SARS-CoV-2, MERS-CoV, HCoV-HKU1, HCoV-OC43, HCoV-NL63 and HCoV-229E. Transfected cells were incubated for 36-48 h at 37°C. Post incubation cells were trypsinized to prepare a single cell suspension and were distributed into 96-well plates. Serum samples were prepared as 3-fold serial titrations in FACS buffer (1x PBS, 2% FBS, 1 mM EDTA), starting at 1:30 dilution, 6 dilutions. 50 μl/well of the diluted samples were added into the cells and incubated on ice for 1h. The plates were washed twice in FACS buffer and stained with 50 μl/well of 1:200 dilution of R-phycoerythrin (PE)-conjugated mouse anti-human IgG Fc antibody (SouthernBiotech #9040-09) and 1:1000 dilution of Zombie-NIR viability dye (BioLegend) on ice in dark for 45min. After another two washes, stained cells were analyzed using flow cytometry (BD Lyrics cytometers), and the binding data were generated by calculating the percent (%) PE-positive cells for antigen binding using FlowJo 10 software. CR3022, a SARS-CoV-1 and SARS-CoV-2 spike binding antibody, and dengue antibody, DEN3, were used as positive and negative controls for the assay, respectively.

### ELISA binding

96-well half-area plates (Corning cat. #3690, Thermo Fisher Scientific) were coated overnight at 4°C with 2 ug/ml of mouse anti-His-tag antibody (Invitrogen cat. #MA1-21315-1MG, Thermo Fisher Scientific) in PBS. Plates were washed 3 times with PBS plus 0.05% Tween20 (PBST) and blocked with 3% (wt/vol) bovine serum albumin (BSA) in PBS for 1 h. After removal of the blocking buffer, the plates were incubated with His-tagged spike proteins at a concentration of 5 ug/ml in 1% BSA plus PBS-T for 1.5 hr at room temperature. After a washing step, perturbed and lotus serum samples were added in 3-fold serial dilutions in 1% BSA/PBS-T starting from 1:30 and 1:40 dilution, respectively, and incubated for 1.5 hr. CR3022 and DEN3 human antibodies were used as a positive and negative control, respectively, and added in 3-fold serial dilutions in 1% BSA/PBS-T starting at 10 ug/ml. After the washes, a secondary antibody conjugated with alkaline phosphatase (AffiniPure goat anti-human IgG Fc fragment specific, Jackson ImmunoResearch Laboratories cat. #109-055-008) diluted 1:1000 in 1% BSA/PBS-T, was added to each well. After 1 h of incubation, the plates were washed and developed using alkaline phosphatase substrate pNPP tablets (Sigma cat. #S0942-200TAB) dissolved in a stain buffer. The absorbance was measured after 8, 20, and 30 minutes, and was recorded at an optical density of 405 nm (OD405) using a VersaMax microplate reader (Molecular Devices), where data were collected using SoftMax software version 5.4. The wells without the addition of serum served as a background control.

### BioLayer Interferometry binding

An Octet K2 system (ForteBio) was used for performing the binding experiments of the coronavirus spike proteins with serum samples. All serum samples were prepared in Octet buffer (PBS plus 0.1% Tween20) as 1:40 dilution, random-biotinylated S proteins were prepared at a concentration of 100nM. The hydrated streptavidin biosensors (ForteBio) first captured the biotinylated spike proteins for 60s, then transferred into Octet buffer for 60s to remove unbound protein and provide the baseline. Then, they were immersed in diluted serum samples for 120s to provide association signal, followed by transferring into Octet buffer to test for disassociation signal for 240s. The data generated was analyzed using the ForteBio Data Analysis software for correction and curve fitting, and for calculating the antibody dissociation rates (k_off_ values) or KD values for monoclonal antibodies.

### Flow cytometry B cell profiling and mAb isolation with HCoV S proteins

Flow cytometry of PBMC samples from convalescent human donors were conducted following methods described previously ^22,32,33^. Frozen human PBMCs were re-suspended in 10 ml RPMI 1640 medium (Thermo Fisher Scientific, #11875085) pre-warmed to 37°C containing 50% fetal bovine serum (FBS). After centrifugation at 400 ⨯ g for 5 minutes, the cells were resuspended in a 5 ml FACS buffer (PBS, 2% FBS, 2mM EDTA) and counted. A mixture of fluorescently labeled antibodies to cell surface markers was prepared, including antibodies specific for the T cell markers CD3(APC Cy7, BD Pharmingen #557757), CD4(APC-Cy7, Biolegend #317418) and CD8(APC-Cy7, BD Pharmingen #557760); B cell markers CD19 (PerCP-Cy5.5, Fisher Scientific #NC9963455), IgG(BV605, BD Pharmingen #563246) and IgM(PE); CD14(APC-Cy7, BD Pharmingen #561384, clone M5E2). The cells were incubated with the antibody mixture for 15 minutes on ice in the dark. The SARS-CoV-2 S protein was conjugated to streptavidin-AF488 (Life Technologies #S11223), the HCoV-HKU1 S protein to streptavidin-BV421 (BD Pharmingen #563259) and the HCoV-NL63 S protein to streptavidin-AF647 (Life Technologies #S21374). Following conjugation, each S protein-probe was added to the Ab-cell mixture and incubated for 30 minutes on ice in the dark. FVS510 Live/Dead stain (Thermo Fisher Scientific, #L34966) in the FACS buffer (1:300) was added to the cells and incubated on ice in the dark for 15 minutes. The stained cells were washed with FACS buffer and re-suspended in 500 μl of FACS buffer/10-20 million cells, passed through a 70 μm mesh cap FACS tube (Fisher Scientific, #08-771-23) and sorted using a Beckman Coulter Astrios sorter, where memory B cells specific to S protein proteins were isolated. In brief, after the gating of lymphocytes (SSC-A vs. FSC-A) and singlets (FSC-H vs. FSC-A), live cells were identified by the negative FVS510 Live/Dead staining phenotype, then antigen-specific memory B cells were distinguished with sequential gating and defined as CD3-, CD4-, CD8-, CD14-, CD19+, IgM-and IgG+. Subsequently, the S protein specific B cells were identified with the phenotype of AF488+BV421+ (SARS-CoV-2/HCoV-HKU1 S protein double positive) or AF488+AF647+ (SARS-CoV-2/HCoV-NL63 S protein double positive). Positive memory B cells were then sorted and collected at single cell density in 96-well plates. Downstream single cell IgG RT-PCR reactions were conducted using Superscript IV Reverse Transcriptase (Thermo Fisher, # 18090050), random hexamers (Gene Link # 26400003), Ig gene-specific primers, dNTP, Igepal, DTT and RNAseOUT (Thermo Fisher # 10777019). cDNA products were then used in nested PCR for heavy/light chain variable region amplification with HotStarTaq Plus DNA Polymerase (QIAGEN # 203643) and specific primer sets described previously ^34,35^. The second round PCR exploited primer sets for adding on the overlapping region with the expression vector, followed by cloning of the amplified variable regions into vectors containing constant regions of IgG1, Ig Kappa, or Ig Lambda using Gibson assembly enzyme mix (New England Biolabs #E2621L) after confirming paired amplified product on 96-well E gel (ThermoFisher #G720801). Gibson assembly products were finally transformed into competent E.coli cells and single colonies were picked for sequencing and analysis on IMGT V-Quest online tool (http://www.imgt.org) as well as downstream plasmid production for antibody expression.

### Neutralization assay

Under BSL2/3 conditions, MLV-gag/pol and MLV-CMV plasmids were co-transfected into HEK293T cells along with full-length or variously truncated SARS-CoV1 and SARS-COV2 spike plasmids using Lipofectamine 2000 to produce single-round of infection competent pseudo-viruses. The medium was changed 16 hours post transfection. The supernatant containing MLV-pseudotyped viral particles was collected 48h post transfection, aliquoted and frozen at −80 °C for neutralization assay. Pseudotyped viral neutralization assay was performed as previously described with minor modification (Modified from TZM-bl assay protocol ^36^). 293T cells were plated in advance overnight with DMEM medium +10% FBS + 1% Pen/Strep + 1% L-glutamine. Transfection was done with Opti-MEM transfection medium (Gibco, 31985) using Lipofectamine 2000. The medium was changed 12 hours after transfection. Supernatants containing the viruses were harvested 48h after transfection. 1) Neutralization assay for plasma. plasma from COVID donors were heat-inactivated at 56°C for 30 minutes. In sterile 96-well half-area plates, 25μl of virus was immediately mixed with 25 μl of serially diluted (3x) plasma starting at 1:10 dilution and incubated for one hour at 37°C to allow for antibody neutralization of the pseudotyped virus. 10,000 HeLa-hACE2 cells/ well (in 50ul of media containing 20μg/ml Dextran) were directly added to the antibody virus mixture. Plates were incubated at 37°C for 42 to 48 h. Following the infection, HeLa-hACE2 cells were lysed using 1x luciferase lysis buffer (25mM Gly-Gly pH 7.8, 15mM MgSO4, 4mM EGTA, 1% Triton X-100). Luciferase intensity was then read on a Luminometer with luciferase substrate according to the manufacturer’s instructions (Promega, PR-E2620). 2) Neutralization assay for monoclonal antibodies. In 96-well half-area plates, 25ul of virus was added to 25ul of five-fold serially diluted mAb (starting concentration of 50ug/ml) and incubated for one hour before adding HeLa-ACE2 cell as mentioned above. Percentage of neutralization was calculated using the following equation: 100 X (1 – (MFI of sample – average MFI of background) / average of MFI of probe alone – average MFI of background)).

### Antibody dependent enhancement assay

Ex vivo antibody dependent enhancement (ADE) quantification was measured using a focus reduction neutralization assay. Monoclonal antibodies were serially diluted in complete RPMI and incubated for 1 hour at 37°C with SARS-CoV-2 strain USA-WA1/2020 (BEI Resources NR-52281) [MOI=.01], in a BSL3 facility. Following the initial incubation, the mAb-virus complex was added in triplicate to 384-well plates seeded with 1E4 of K562 or Daudi cells and were incubated at 34°C for 24 hours. 20µL of the supernatant was transferred to a 384-well plate seeded with 2E3 HeLa-ACE2 cells and incubated for an additional 24 hours at 34°C. Plates were fixed with 25 ul of 8% formaldehyde for 1 hour at 34°C. Plates were washed 3 times with 1xPBS 0.05% Tween-20 following fixation. 10µL of human polyclonal sera diluted 1:500 in Perm/Wash Buffer (BD Biosciences)was added to the plate and incubated at RT for 2 hours. The plates were then washed 3 times with 1xPBS 0.05% Tween-20 and stained with peroxidase goat anti-human Fab (Jackson Scientific, 109-035-006) diluted 1:2000 in Perm/wash buffer then incubated at RT for 2 hours. The plates were then washed 3 times with 1xPBS 0.05% Tween-20. 10µL of Perm/Wash buffer was added to the plate then incubated for 15 minutes at RT. The Perm/Wash buffer was removed and 10µL of TrueBlue peroxidase substrate (KPL) was added. The plates were incubated for 30 minutes at RT then washed once with milli-Q water. The FFU per well was then quantified using a compound microscope. The PFU/mL of the monocyte plate supernatant was calculated and graphed using Prism 8 software.

### Negative Stain Electron Microscopy

The HCoV-HKU1 S protein was incubated with a 3-fold molar excess of Fab CC40.8 for 30 mins at room temperature and diluted to 0.03 mg/ml in 1X TBS pH 7.4. 3 μL of the diluted sample was deposited on a glow discharged copper mesh grid, blotted off, and stained for 55 seconds with 2% uranyl formate. Proper stain thickness and particle density was assessed on a FEI Morgagni (80keV). The Leginon software ^37^ was used to automate data collection on a FEI Tecnai Spirit (120keV), paired a FEI Eagle 4k x 4k camera. The following parameters were used: 52,000x magnification, −1.5 μm defocus, a pixel size of 2.06 Å, and a dose of 25 e^−^/Å^2^. Micrographs were stored in the Appion database ^38^, particles were picked using DogPicker ^39^, and a particle stack of 256 pixels was made. RELION 3.0 ^40^ was used to generate the 2D class averages. The flexibility of the fab relative to the spike precluded 3D reconstruction.

### Statistical Analysis

Statistical analysis was performed using Graph Pad Prism 8 for Mac, Graph Pad Software, San Diego, California, USA. Median area-under-the-curve (AUC) or reciprocal 50% binding (ID50) or neutralization (IC50) titers were compared using the non-parametric unpaired Mann-Whitney-U test. The correlation between two groups was determined by Spearman rank test. Data were considered statistically significant at * p < 0.05, ** p < 0.01, *** p < 0.001, and **** p < 0.0001.

## Data availability

The authors declare that the data supporting the findings of this study are available within the paper and its supplementary information files or from the corresponding author upon reasonable request. Antibody sequences have been deposited in GenBank under accession numbers XXX-XXX. Antibody plasmids are available from Dennis Burton under an MTA from The Scripps Research Institute.

